# D^2^BGAN: Dual Discriminator Bayesian Generative Adversarial Network for Deformable MR-Ultrasound Registration Applied to Brain Shift compensation

**DOI:** 10.1101/2022.01.22.477329

**Authors:** M. Rahmani, H. Moghadassi, P. Farnia, A. Ahmadian

## Abstract

**Purpose:** In neurosurgery, image guidance is provided based on the patient to pre-operative data registration with a neuronavigation system. However, the brain shift phenomena invalidate the accuracy of the navigation system during neurosurgery. One of the most common approaches for brain shift compensation is using intra-operative ultrasound (iUS) imaging followed by registration of iUS with pre-operative magnetic resonance (MR) images. While, due to the unpredictable nature of brain deformation and the low quality of ultrasound images, finding a satisfactory multimodal image registration approach remains a challenging task.

**Methods:** We proposed a new automatic unsupervised end-to-end MR-iUS registration approach based on the Dual Discriminator Bayesian Generative Adversarial Network (D^2^BGAN). The proposed network consists of two discriminators and is optimized by introducing a Bayesian loss function to improve the generator functionality and adding a mutual information loss function to the discriminator for similarity measurement. An evaluation was performed using the RESECT training dataset based on the organizer’s manual landmarks.

**Results:** The mean Target Registration Error (mTRE) after MR-iUS registration using D^2^BGAN reached 0.75±0.3 mm. The D^2^BGAN illustrated a clear advantage by 85% improvement in the mTRE of MR-iUS registration over the initial error. Also, the results confirmed that the proposed Bayesian loss function rather than the typical loss function outperforms the accuracy of MR-iUS registration by 23%.

**Conclusion:** The D^2^BGAN improved the registration accuracy while allowing us to maintain the intensity and anatomical information of the input images in the registration process. It promotes the advancement of deep learning-based multi-modality registration techniques.

## I. INTRODUCTION

Extensive tumor removal is critical to the success of surgical tumor excision and increases the survival rate of patients, especially in neurosurgery. Maximal safe resection of brain tumors while preserving normal tissue is a challenging task due to its close resemblance to normal brain parenchyma in addition to the existence of vital structures in neighboring areas [8, 19]. Recently, neuronavigation technologies have inspired tremendous capacity in providing accurate localization of the tumors and vital structures of the brain during surgery. These systems update pre-operative images of patients such as Magnetic Resonance (MR) images and Computed Tomography via registration of patient coordinate achieved by the tracked tool with pre-operative data [7]. However, neuronavigation systems could not serve an accurate guide due to intra-operative brain deformation, known as brain shift [2]. Brain shift phenomena are caused by a number of physical, biological, and surgical factors in cortical and deep brain regions and invalidate pre-operative data [19].

On the other hand, intra-operative Ultrasound (iUS) is found as a promising imaging technique for improving the perception of anatomical structure and compensation for brain shift during neurosurgery and has acquired a lot of traction, recently [17]. The iUS updates patients’ coordination during neurosurgery via registration of the iUS with pre-operative MR images. Therefore, MR-iUS registration is needed to implement during surgery, which is a challenging task due to the low quality and lack of detail in the ultrasound images as well as the complex and unpredictable nature of brain deformation [12]. Numerous investigations have attempted to overcome the constraints of multimodal image registration methods for brain shift compensation including feature-based [22] and intensity-based [30] methods. Unlike feature-based methods, which rely on feature selection accuracy, intensity-based methods appeared to be more successful in this area [6]. For this purpose, Rivas et al. [24] presented a novel algorithm for MR-iUS Registration via PaTch-based cOrrelation Ratio (RaPTOR), which computes the local correlation ratio. Farnia et al. [4, 5] based on improving the residual complexity value in the wavelet and curvelet domains, suggested hybrid techniques for aligning echogenic structures in MR with iUS images. Zhou and Rivaz [35] presented a non-rigid symmetric registration approach based on pre- and post-resection ultrasound images for brain shift compensation and residual tumor assessment that is difficult to evaluate on typical post-resection images. Riva et al. [23] evaluated the brain shift compensation at three time points during surgery. The first point was before dural opening with a linear correlation of linear combination, after dural opening with normalized cross-correlation registration and deformable B-spline registration methods. Finally, after brain tumor resection, iUS was applied to the pre-operative image. Wein proposed a model based on the LC^2^ multi-modal similarity measure [29, 30]. Drobny et al. [3] presented a block matching approach using NiftyReg [18] for automatic MR-iUS registration.

In the CuRIOUS 2018 [32], the first independent Challenge of MR-iUS registration algorithms, Machado et al. [16] proposed a method for Deformable Registration via Attribute Matching and Mutual-Saliency Weighting (DRAMMS) algorithm [13] for the MR-iUS registration. Heinrich et al. [11] implemented Deeds/SSC, which comprises linear and non-rigid registration, both based on discrete optimization and modality-invariant image features. However, the development of these algorithms has led to an increase in registration accuracy but increased the computational complexity of the registration process, which is not suitable for real-time compensation of brain shift.

On the other hand, Deep Learning (DL) has recently shown great potential and gained attention in medical applications due to its superior performance in a variety of image processing applications, such as object detection [20], feature extraction [28], image segmentation [1], etc. For the first time, DL approaches were used for brain shift correction in CuRIOUS 2018. Sun et al. [27] presented a DL framework for non-rigid MR-iUS registration using a 3D Convolutional Neural Network (CNN) consisting of a feature extractor, deformation field generator, and a spatial sampler. As an imitation game, Zhong et al. [34] suggested a learning-based strategy for intra-operative brain shift correction. They trained a neural network to imitate the demonstrator’s behavior and anticipate the optimal deformation vector. Zeineldin et al. [33] used 3D CNN for deformable MR-iUS image registration.

Automated DL approaches for MR-ultrasound registration reduce the registration time by learning a global function for multi-modal non-rigid image registration, which can be performed quickly on a pair of test images. Therefore, all of these methods are suitable for real-time compensation of brain shift, but they were not able to match the images accurately enough. Therefore, achieving high accuracy along with high speed of registration is a major issue that remains a challenging task for brain shift compensation during neurosurgery. Furthermore, the non-rigid and unpredictable nature of brain deformation led us to unsupervised learning in which the challenges associated with the ground truth data generation and optimization methods would be eliminated.

To address the multi-modal image registration challenges with the benefit of DL methods, in this study for the first time, we propose an unsupervised end-to-end registration method via Dual Discriminator Bayesian Generative Adversarial Network (D^2^BGAN). The presented network consists of two kinds of neural networks based on deep convolutional Generative Adversarial Network (GAN), a generator, and two discriminators [21] and dual discriminator GAN [14]. We optimize the unsupervised end-to-end architecture of the dual discriminator GAN by introducing a Bayesian loss function to improve the generator functionality and add a mutual information loss function to the discriminator.

## II. MATERIALS AND METHODS

In this section, in addition to the dataset and basic theory of GAN, we would propose our registration approach and define the design of loss functions we have presented in the proposed method.

### A. Dataset

In this study, experiments were conducted on the public RESECT dataset [31], which contains pre-operative MR and iUS images of 22 patients with low-grade glioma who have received surgeries at St. Olavs University Hospital, Norway.

Each patient was scanned using Gadolinium-enhanced T1-weighted and T2-FLAIR MR protocols with the 1.5T Siemens Magnetom Avanto to reveal the anatomy and pathology with a voxel size of 1×1×1 mm^3^ and a resolution of 256×256×192 pixels. The series of B-mode US images were obtained with a 12FLA-L linear probe with a voxel size of 0.14 × 0.14 × 0.14 mm^3^ to 0.24 × 0.24 × 0.24 mm^3^ before, during, and after tumor resection to track the surgical progress and tissue deformation. Overall, 338 corresponding anatomical landmarks were identified between MR and US. 264 paired slices were extracted from the 3D volume of MR and US. Each network was trained on an Intel^®^ Core ™ i9-9900k CPU @ 3.60GHz with 32.0GB RAM and a GeForce RTX 3060Ti.

### B. Generative Adversarial Network (GAN)

The rise of GANs [10] has given medical image processing a whole new era. A GAN, in particular, does not need labeled data to attain an exquisite outcome, which could be generated through the Generator (G) and Discriminator (D) networks competition. As a result, GANs are fast proving to be a cutting-edge basis, achieving better results in various medical applications including image registration [9].

GAN achieves stepped forward performance by permitting two models to be trained simultaneously. They will learn in an adversarial manner; hence the G and D models are used to represent data distribution and estimation of samples, respectively. G attempted to produce different approaches to understand discriminative potential by simulating real data and providing it with the ability to verify its authenticity.

In the GAN optimization process, G tries to find an optimal solution in which D recognizes real data erroneously. Therefore, to address this issue and estimate the real or fake data probability, the objective function for D can be defined as:

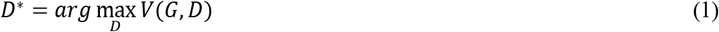

where V (G, D) is formulated as follows:

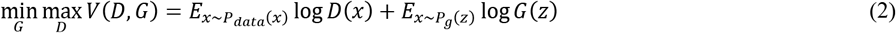

where x and z are the real data and noise in the input of G, respectively. G(z) is the image generated by G, and D(x) is the probability to find out the real data. G will learn from real and fake data with a probability the distribution *P_data_*(*x*) from the data x and *P_g_*(*z*) of the input noise. Consequently, the optimization formulation of G can be written as:

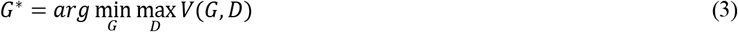

### C. D^2^BGAN framework

The proposed procedure of the registration in the *D^2^BGAN* pipeline with two discriminators is shown in Fig. 1.

**Fig. 1.**
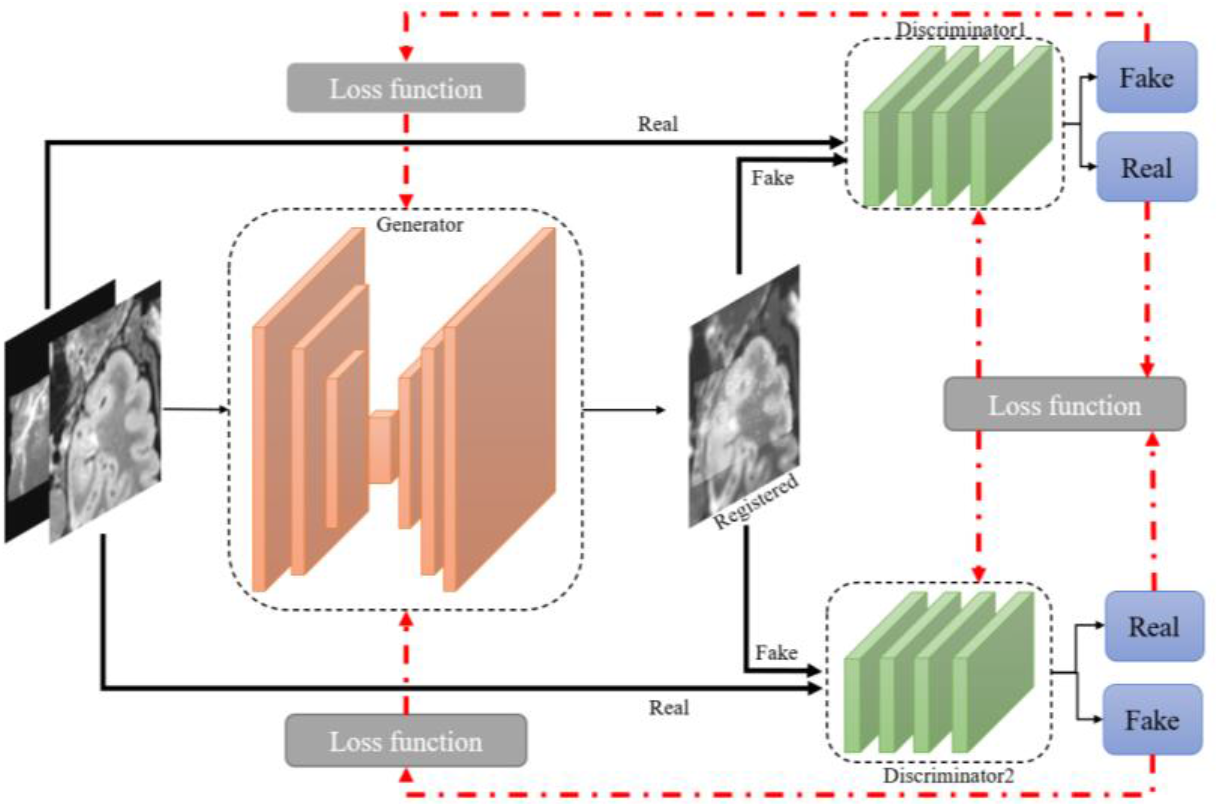
D2BGAN framework.

Since G is trained with the inputs, the registered images are inspired to be realistic with the detail of input images to fool the discriminators D1 and D2. Based on the similarity, G is the architect of a CNN, which allows the network to predict in a dense pixel-wise manner. High-resolution activation maps are integrated with the up-sampled outputs and delivered to the convolution layers to gather more accurate outputs and better localization. Therefore, the generator is an autoencoder that consists of an encoder and decoder network, as illustrated in Fig. 2.

**Fig. 2.**
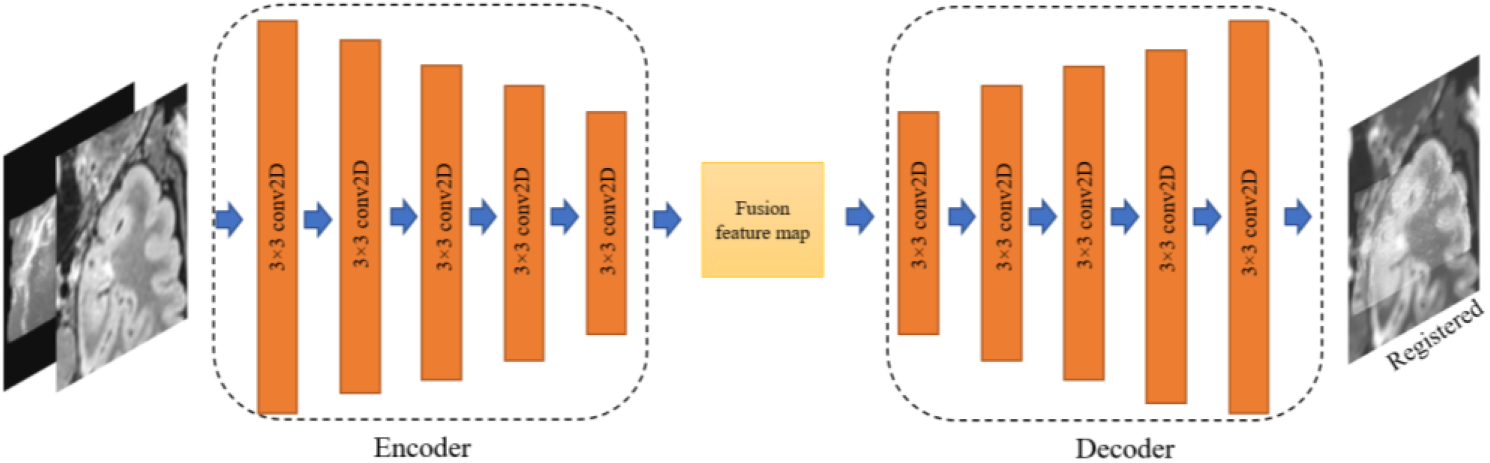
The architecture of the generator network with 3 × 3: filter size, Conv2D: convolutional layer which obtains k feature maps, with batch normalization and ReLU active function.

The encoder takes the input images and performs feature extraction and fusion, which the output of the encoder is a fusion feature map state. The encoded state is fed to the decoder for registered image reconstruction. The encoder consists of 5 convolutional layers and each layer can obtain by 3 × 3 filter size. The decoder is a four-layer network as illustrated in Fig. 2. To avoid exploding/vanishing gradients and speed up the training process, batch normalization is applied to all convolutional layers and the strides of all layers are set as 2. In addition, the ReLU activation function is used to accelerate convergence.

D1 and D2 are trained to distinguish between MR-iUS images with the registered image, respectively. Therefore, discriminators would be a two-channel layer having both the sampled data and the source image. In a dual discriminator structure, in addition to maintaining balance between discriminators should also consider the adversarial relationship between the generator and the discriminators. Since in the training process, the strength or weakness of each discriminator will eventually lead to the inefficiency of the other.

The design of the network and training procedures are used to attain balance in this architecture by having the same architecture, as shown in Fig. 3. All convolutional layers have a stride of 2, with *ReLU* active function and batch normalization. The last layer is a dense layer with a *Tanh* activation function to predict the likelihood of the input picture from source images instead of G.

**Fig. 3.**
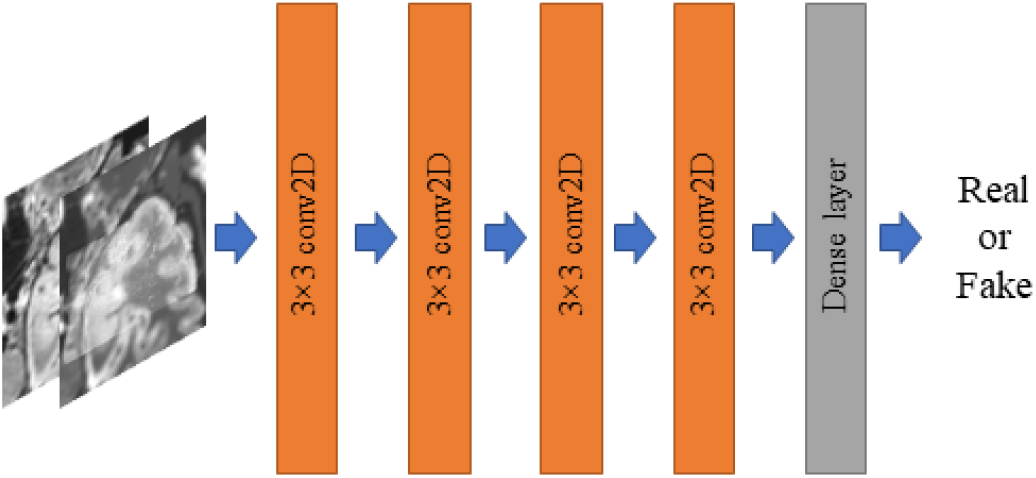
The overall architecture of our discriminator. Conv2D convolutional layer, which obtains feature maps with 3 × 3 filter size, batch normalization, and dense layer.

G will begin to generate images randomly when the generator is unable to deceive the D with adversarial relationship establishment fail, which would distort the generative model. Therefore, the training target of G is defined by minimizing the adversarial objective, as follow [25]:

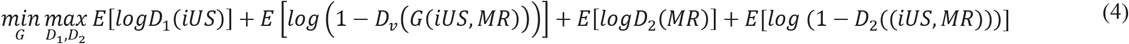

which the discriminators’ goal is to maximize Eq. (4). The divergence between the probability distributions of the generator G and the two real discriminators, D1 and D2, will become smaller.

In the following, a measure of distance between probability distributions is used to define the G and D losses. The fake data distribution can only be affected by the G, therefore, during generator training, we drop the real data. The D would experience the inverse of the same scenario. Even though they are derived from the same formula, the G and D losses are different in the end. Therefore, the G loss function as an adversarial loss *L_G_* is defined as:

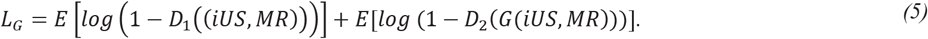

The discriminators are responsible for distinguishing between source and moving images with the registered image. The Jensen-Shannon divergence between distributions can be calculated using adversarial losses of discriminators, which can be used to determine if the intensity or texture information is unrealistic. As a result, matching the realistic distribution is encouraged. The following formulas are the definition of adversarial losses:

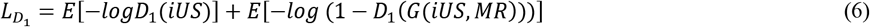

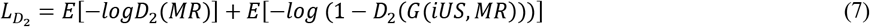

To improve network performance, two other loss functions, one for implementation in G and the other for D, are presented as follow:

#### i. Mutual information loss function

The quantity of information learned about the registered image from the other input images is called Mutual Information (MI). Therefore, we’re looking for mutual dependence between source and fixed images with the registered image. MI for images X and Y is defined as:

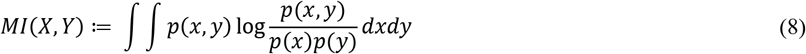

Where *p*(*x*) and *p*(*y*) are the marginal probability densities and *p*(*x, y*) is the joint probability density of *X* and *Y* [26]. The *D*_1_ and *D*_2_ loss function could be defined based on the MI of the registered and the input images, MR and iUS, therefore, we would define MI loss function for *D*_1_ and *D*_2_ as:

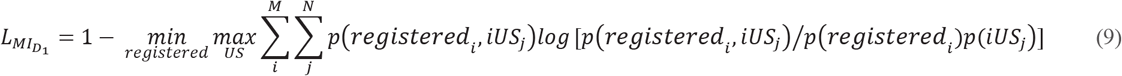

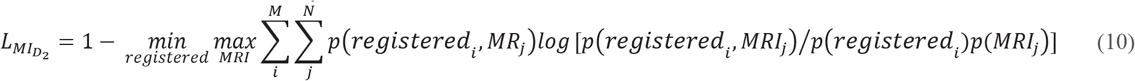

As a result of the MI loss, the discriminator model is regularized and implemented in the training and image generating process.

#### ii. Bayesian loss function

Based on the prior knowledge of MR and iUS images, Bayes’ theorem defines the likelihood of the registered image with the input images. The Bayesian theory is an interpretation of the probability concept, rather than image intensity and similarity. It should be considered a reasonable expectation representing a state of knowledge.

The independency of the Bayes’ theorem on the similarity between pixels’ intensity makes it very efficient and instructive for this nonlinear problem that is pixel intensity independent. The notion of conditional probability is used to derive Bayes’ theorem:

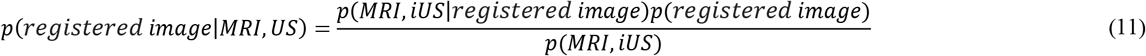

where *P(registered image)* and *P(MRI,iUS)* are the known prior probabilities and *p(MRI, iUS\registered image)* is known as conditional probability, in which M and N are the numbers of pixels in the registered image and input images, respectively.

One advantage of Bayesian probability is with the determined posterior distribution which would make the computation simple. Bayesian approaches enable the prediction of the likelihood of the registered image, which is referred to as the predictive probability. The predictive distribution represents the probability distribution of the registered image based on the MR and US images. Therefore, to the best of our knowledge, for the first time, based on the Bayes’ theorem, we would define the Bayes’ loss function for the generator as:

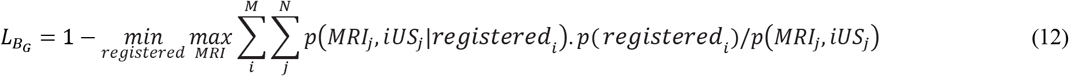

### D. Evaluation Metric

RESECT dataset includes corresponding landmarks in the MR and iUS images which are annotated by an expert as mentioned in section II. We used the common evaluation metric of the mean Target Registration Error (mTRE) for MR-iUS registration, similar to others in this field [15, 32], which is the average distance between corresponding landmarks in each pair of MR and US images. The mTRE is defined as:

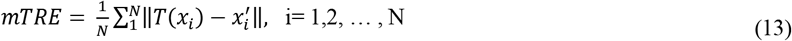

where *x_i_* and 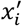 represent corresponding landmark locations annotated by an expert in the MR and iUS images, respectively, and T would be the MR to iUS transform calculated from the registration procedure.

## III. RESULTS

To evaluate the performance of the unsupervised end-to-end D^2^BGAN registration method for brain-shift compensation, we compared the mTRE of identifying anatomical landmarks between MR and iUS in registered images. Table 1 summarizes the mTREs of pre-and post-registration of the proposed method for all the trained 22 RESECT cases, individually. The multimodal registration was performed using two different sets of loss functions including *L*_1_ and *L*_2_. The *L*_1_ loss function is used *L_G_*, *L*_*d*1_ and *L*_*d*2_ and *L_G_* + *L_B_G__*, *L*_*D*_1__ + *L*_*MI*_*D*_1___ and *L*_*D*_2__ + *L*_MI_D1__ is used in *L_2_* loss function.

**Table 1.** mTRE of unsupervised end-to-end D^2^BGAN approach evaluated with corresponding landmarks.

| Cases | mTRE (mm):<br>Before Registration | mTRE (mm): After Registration |  |
| --- | --- | --- | --- |
| | | $L_1$ loss function | $L_2$ loss function |
| #1 | 1.86 | 0.8184 | 0.7377 |
| #2 | 5.75 | 1.7625 | 1.1266 |
| #3 | 9.63 | 1.5289 | 0.9664 |
| #4 | 2.98 | 0.8520 | 0.5183 |
| #5 | 12.20 | 1.5311 | 1.4715 |
| #6 | 3.34 | 0.4644 | 0.2992 |
| #7 | 1.88 | 0.7826 | 0.5380 |
| #8 | 2.65 | 0.7109 | 0.6584 |
| #12 | 19.76 | 1.2702 | 0.8793 |
| #13 | 4.71 | 0.59665 | 0.5946 |
| #14 | 3.03 | 0.5196 | 0.4340 |
| #15 | 3.37 | 1.0292 | 0.8440 |
| #16 | 3.41 | 1.1579 | 1.1256 |
| #17 | 6.41 | 0.6660 | 0.8418 |
| #18 | 3.66 | 1.2277 | 0.7604 |
| #19 | 3.16 | 1.2741 | 1.0548 |
| #21 | 4.46 | 1.5884 | 0.7350 |
| #23 | 7.05 | 0.7250 | 0.4869 |
| #24 | 1.13 | 0.4679 | 0.3606 |
| #25 | 10.10 | 1.0692 | 0.8131 |
| #26 | 2.93 | 0.4731 | 0.3179 |
| #27 | 5.86 | 1.0950 | 1.0493 |
| Mean ± std | 5.4240± 4.2901 | 0.9823±0.3958 | 0.7551±0.3017 |

A clear advantage over the initial error can be seen from 5.42 mm to 0.75 mm. Furthermore, the variations of mTRE for 22 case is depicted using distribution plots in Fig. 4. The y-axis shows a box plot with the mTRE for each combination (maximum, minimum, interquartile range, and median).

**Fig. 4.**
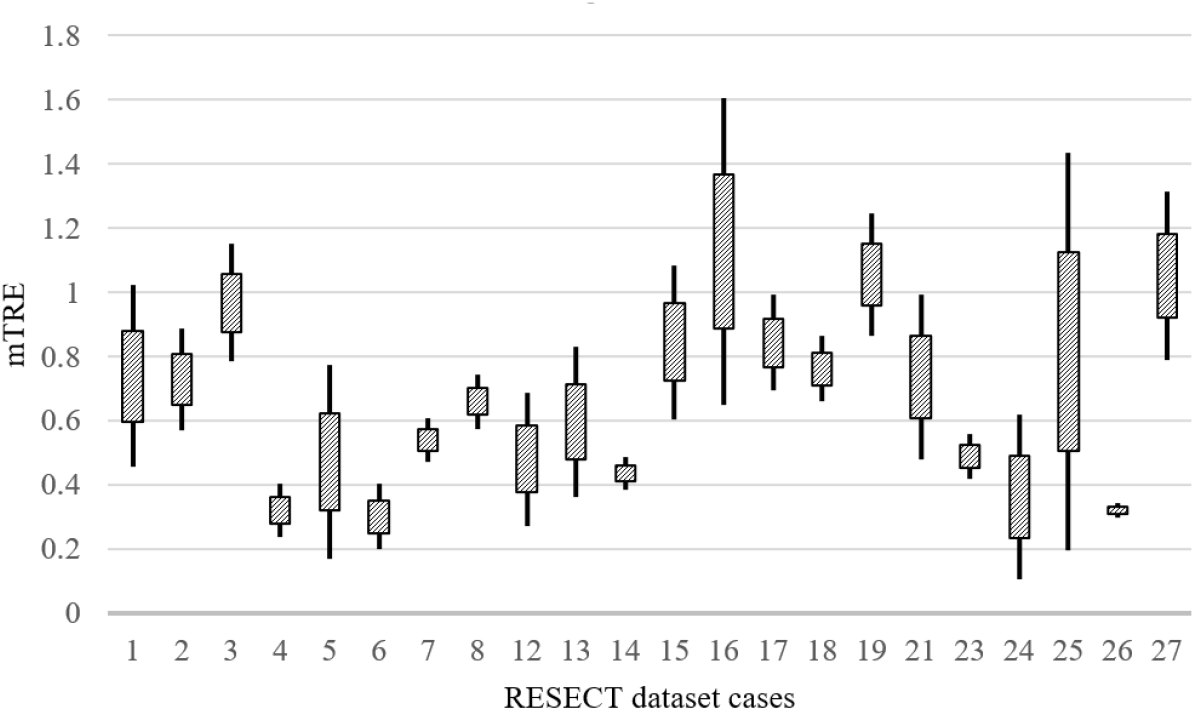
Distribution plots of mTRE for 22 cases.

To evaluate registration results, qualitatively, Fig. 5 provides visual examples of MR to iUS registration for cases 9, 15, and 18 of datasets. The last row depicts a color overlay iUS on top of the pre-operative MR which demonstrates the alignment has been improved.

**Fig. 5.**
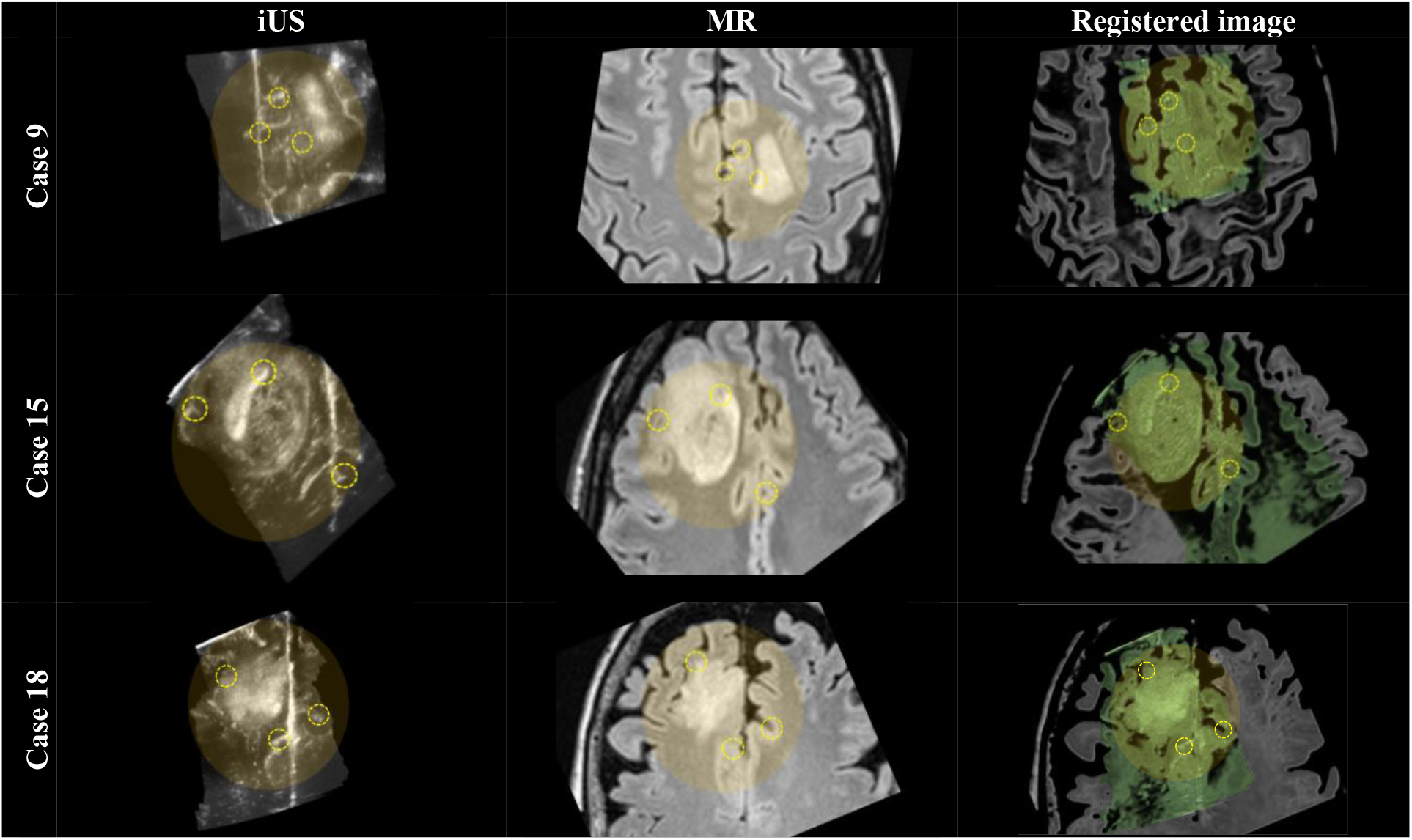
MR-to-iUS registration results. The first, second, and third rows correspond to cases 9, 15, and 18, respectively. The left, middle, and right columns correspond to the iUS and pre-operative MR, and the registered images, respectively. In the yellow shadow circles, corresponding structures are illustrated with dashed yellow circles.

Also, Fig. 6 indicates the brain shift in sagittal, axial, and coronal slices. *Rows* indicate sequentially pre-operative imaging MR, iUS, and registered images. The overlay between the iUS acquisition and the pre-operative MR is shown in the last row.

**Fig. 6.**
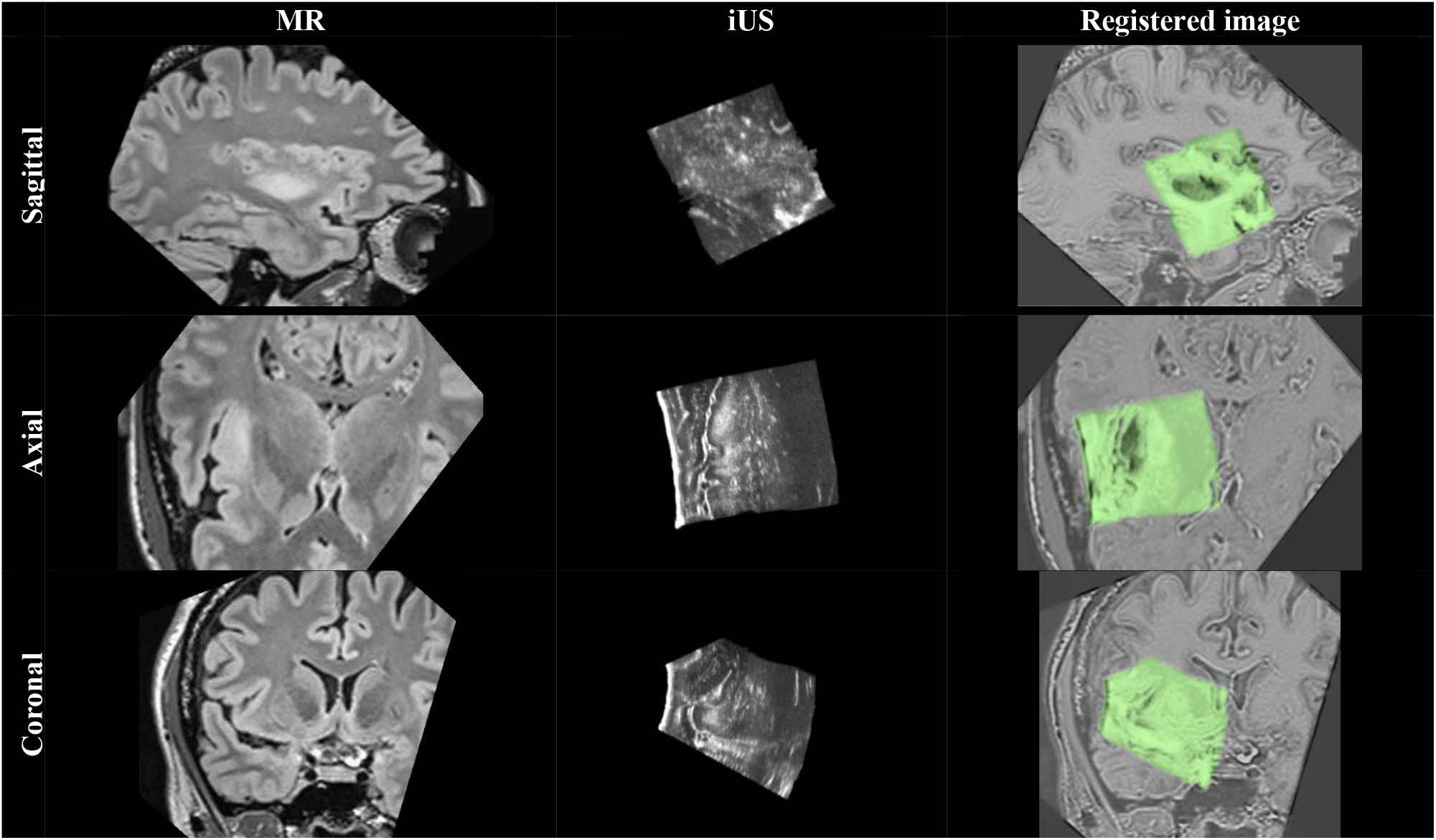
the brain-shift indicate in sagittal, axial, and coronal planes of case 4 in RESECT dataset. *Rows* indicate sequentially pre-operative MR, iUS, and registered images.

In order to compare our proposed method to others, Table. 2 presents final landmarks errors for the proposed D^2^BGAN and other methodologies found in the literature for MR-iUS registration, performed on the RESECT database. Therefore, a Comparison of our registration methods against other MR-iUS registration approaches such as LC^2^ [30], SSC/Deeds [11], NiftyReg [18], cDRAMMS [13] as well as learning studies, FAX [34], CNN [27] is done.

**Table 2.** mTRE for our proposed methods and the state-of-the-art methods on the RESECT dataset.

| method | mTRE (mean±std) |
| --- | --- |
| LC2 | 1.75 ± 0.62mm |
| SSC | 1.67±0.54 mm |
| NiftyReg | 2.90 ±3.59 mm |
| cDRAMMS | 2.28±0.71 mm |
| FAX | 1.21±0.55 mm |
| CNN | 3.91± 0.53mm |
| D<sup>2</sup>BGAN | 0.75± 0.30mm |

## IV. DISCUSSION

We proposed a new automatic DL-based MR-iUS registration approach by constructing a D^2^BGAN that is trained in an unsupervised end-to-end manner, applied in image-guided neurosurgery. The presented approach would eliminate the dependence of image registration in pixel intensities variation in different modalities so that the network has the ability to learn non-linear transformations.

All experiments were run on the all RESECT training scans dataset and evaluation have done using all manual landmarks which have been produced by the organizers. No manual initialization is required in this fully automated algorithm. The network has been optimized by introducing a Bayesian loss function to improve the generator functionality and also adding a mutual information loss function to the discriminator. The Bayesian loss function would eliminate the dependence of image registration in pixel intensities variation of different modalities and the similarity measure using the mutual information loss function has increased the network capability. As shown in Table 1, a significant improvement is revealed in the results over the initial alignment. The D^2^BGAN network with the L_1_ loss function reduced the initial mTRE by about 81%. This is while changing it to the proposed L_2_ loss function, the mTRE reduction is about 86%. By changing the loss function from L_1_ to L_2_, in cases 5,13,16, and 27 the error rate has not changed, but in some cases such as 6 and 23, we had the most changes, about 65% rather than the initial alignment.

The proposed algorithm, in addition to increasing the accuracy of MR-iUS registration, would reduce its time which is a remarkable advantage due to the importance time in neurosurgery. This would allow us to provide the surgeon with real-time image registration. Besides, the algorithm must be capable of registering different fields of view in multi-modality registration. It can be seen in Fig. 5 that shows the result of the MR-iUS registration in the sagittal, axial, and coronal plane for case 4 of the RESECT dataset.

Particularly, the D^2^BGAN inclusion as a registration method had a significant influence on registration accuracy and has given us the capability to preserve the intensity and anatomical information. The network performance has enabled us to have an intensity and scale-invariant transform with a great extent in the registered image which demonstrates the efficiency of utilizing it for multi-modal registration. Although there is no relationship between the pixel intensities in the two image modalities, this does not mean that the nature of the images is lost in the registration process. The results indicated that the D^2^BGAN can conduct automatic accurate deformable MR-iUS image registration, and as a result, it could be implemented in image-guided neurosurgical interventions.

## V. CONCLUSION

Neuronavigation technology enables guidance and finds the best route to the target in neurosurgical procedures. However, neuronavigation systems have been unreliable based on the brain shift that happens during neurosurgery. In order to deal with this phenomenon, MR-iUS registration is provided for brain shift compensation. However, due to the unpredictable nature of brain deformation and the low quality of ultrasound images, finding a satisfactory multimodal registration approach remains a challenging task in this era. Therefore, this study attempted to increase the accuracy of MR-iUS registration to identify intra-operative brain shift via D^2^BGAN.

The D^2^BGAN inclusion as a registration method had a significant influence on registration accuracy and has given us the capability to preserve the intensity and anatomical information. The network performance has enabled us to have an intensity and scale-invariant transform with a great extent in the registered image which demonstrates the efficacy of utilizing it for multi-modal registration. The results of our proposed D^2^BGAN compared to other methods in MR-iUS registration have revealed that the presented method, in addition to real-time MR-iUS registration, also increases the registration accuracy. It encourages the further development of DL-based approaches for multi-modality non-rigid registration.

## STATEMENTS AND DECLARATIONS

### Ethical approval

This article does not contain any studies with human participants or animals performed by any of the authors.

### Informed consent

Informed consent was obtained from all individual participants included in the study.

